# Actomyosin active torques determine body plan handedness in *C. elegans*

**DOI:** 10.64898/2026.02.19.706789

**Authors:** Arittri Mallick, Julia Pfanzelter, Lokesh G. Pimpale, Stephan W. Grill

## Abstract

Left–right (LR) asymmetries are a defining feature of bilaterian body plans, but the mechanisms that convert molecular chirality into a conserved body-axis handedness remain unclear. In *C. elegans*, LR axis specification proceeds via a chiral spindle skew that establishes a handed cell–cell contact pattern (CCP) at the 6-cell stage; an arrangement that, when reversed, leads to *situs inversus* nematodes [1]. Chiral acto-myosin flows that arise from active chiral torques have been previously identified as the driver of this LR asymmetric spindle skew [2]. Here, we show that high amounts of Lifeact::mKate2, widely used to visualize F-actin, reverse the handedness of active torques in the cortex. We find that this results in reversed chiral acto-myosin flows during LR symmetry breaking, leading to mirrored 6-cell embryos and *situs inversus* nematodes. Our findings demonstrate that reversing the handedness of active torques changes the LR handedness at the organismal level and establish active chiral torques as instructive for LR axis specification in worms.

## Introduction

The body plan of bilaterians, which amount to more than 99% of all animals [3], is characterized by three main body axes: the anterior-posterior (AP) axis that denotes head and tail, the dorsal-ventral (DV) axis that denotes back and front, and the left-right (LR) body axis [4]. Establishment of these mutually-orthogonal axes occurs during early development. A common strategy to consistently break LR symmetry on the organismal level is to deploy chiral molecular components [2, 5–12]. Cytoskeletal filaments such as microtubules and actin are often involved in LR symmetry breaking, as they form chiral structures at the microscale. In mouse, frog and fish the beating of microtuble-based cilia induces a chiral, directed flow of extracellular fluid that initiates a signaling cascade that systematically breaks LR symmetry [13–15]. In many other species, both vertebrates and invertebrates, the acto-myosin network is involved in LR symmetry breaking. In Drosophila the motor protein Myo1D is central and mutations in it reverse chirality on cellular and tissue level [7, 8]. In snails mutations in the formin Ldia2 result in animals with a reversed body plan [5, 6]. Both in quails and in *C. elegans* myosin-driven chiral flows participate in the establishment of the LR body axis [2, 12, 16–18]. In quails the chiral flow is on a tissue scale, since many cells rotate around a central node structure [17, 18]. In *C. elegans* chiral flows occur on a subcellular scale, as microscopic torques drive chiral flows in the acto-myosin cortex of the developing embryo [2, 12, 16, 19, 20]. These counter-rotational flows skew the mitotic spindle and hence the division axis of the ABa and ABp cell during the 4-6 cell stage [2, 16]. This results in a handed cell-cell contact pattern (CCP) at the 6-cell stage the embryo [1, 21]. This asymmetry in cell contacts between the four daughter cells enables LR asymmetric cell signaling and the establishment of distinct cell fates on the left and right body halves thereby determining the handedness of the body plan [1, 21–25]. Bill Wood could could show in 1991 that reversing the CCP by pushing the ABa cell with a micro-needle during division is sufficient to reverse the body plan [1]. Chiral flows can result from active torque generation at the molecular scale [2, 16, 20], and we here ask if changing the handedness of active torques can result in an inversion the handedness of the *C. elegans* body plan.

## Results

### Lifeact reverses chiral acto-myosin flows during polarity onset in *C. elegans* zygotes

Stabilization of actin filaments have been shown to reverse chirality in fibroblasts [11]. The expression of Lifeact, a peptide used to label filamentous actin, is known to stabilize F-actin and impact cortical dynamics[26–32]. Previous work from the lab indicated that Lifeact can change chiral actomyosin dynamics in the early *C. elegans* embryo [33]. However, an account of reversal of chirality upon expression of Lifeact has not yet been reported. Here, we discovered that high amounts of Lifeact::mKate2 (SI Fig.1d) can reverse the chirality of cortical flows during polarity onset in the zygote of *C. elegans* (Fig.1a, Movie S1). We demonstrate this by imaging and quantifying cortical flows during polarity establishment in two transgenic worm strains referred to as Lifeact- and Lifeact++ ([2, 19, 34], see Methods). Both strains express endogenously labelled non-muscle myosin II (NMY-2::GFP), while the Lifeact++ strain, in addition, expresses high amounts of the Lifeact peptide (Lifeact::mKate2). We find that while the component of cortical flows along the A-P direction (*v*_x_) during polarity onset is similar between the Lifeact- and Lifeact++ zygotes (SI, Fig.1a, error bars denote SEM), the flow velocity component orthogonal to the A-P direction (*v*_y_) has opposite sign (see Fig.1b, SI Fig.1a, Movie S1). This reversal of flow handedness is reflected in the distributions of instantaneous chiral velocities 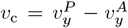, where 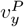 and 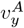 denote the *y*-component of space-averaged instantaneous velocities in the anterior (A) and posterior (P) domains of the zygotes [2] (Fig.1c, see Methods). The *v*_c_ distributions shift from negative values for the Lifeact-condition to positive values for the Lifeact++ condition. Averaging the instantaneous chiral velocities *v*_c_ over time for each embryo (denoted by 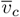), followed by an average over embryos reveals that the average chiral velocity 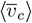 for the ensemble changes sign for Lifeact++ (1.58 ± 1.37 *µm/min*; mean ± error of the mean at 99% confidence, unless otherwise noted) as compared with Lifeact-(−1.55 ± 1.03 *µm/min*). Notably, 24 hours of *mKate2(RNAi)*, which targets Lifeact, rescues handedness reversal in Lifeact++ zygotes 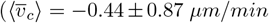 see Fig.1d, Movie S2). We conclude that the presence of high amounts of Lifeact::mKate2 results in cortical flow fields with reversed handedness during polarity establishment.

We next asked if the reversed flow handedness results from a reversal of the handedness of active torques [2, 12, 16, 20]. We used active chiral fluid theory to determine the chirality index *c* of the actomyosin cortex in the zygote, defined as the ratio of active torque density *τ* to active tension *T* [2, 20] (see Methods). We find that *c* = − 0.24(− 0.49, − 0.09) (bootstrapped mean with 95% confidence interval unless otherwise noted; see Methods) for Lifeact++ zygotes, compared with *c* = 0.23(0.10, 0.36) for the Lifeact-condition. We conclude the reversal of the handedness of cortical flow in the Lifeact++ condition likely results from a reversal of the handedness of active torques.

RhoA affects active torque generation in *C. elegans*, and RNAi of its GAP (GTPase Activating Protein) RGA-3 [35– 39] increases chiral flow without affecting handedness [2, 12]. We speculate that increasing RhoA activity in Lifeact++ zygotes should lead to an increase in the magnitude of chiral flows without impacting handedness. Indeed, we find that upon *rga-3(RNAi)* in Lifeact- and Lifeact++ worms (see Methods), both the average chiral velocity 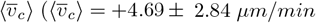 for Lifeact++ and −11.45 ± 4.01 *µm/min* for Lifeact-) and the chirality index c (*c* = −1.30(− 3.17, −0.36) for Lifeact:++; 6.32(2.41, 13.31) for Lifeact-) increase. However, handedness is maintained: Lifeact-undergoes cortical flows with the stereotypic handedness, while Lifeact++ undergoes cortical flows with reversed handedness, as observed in the absence of the *rga-3(RNAi)* (Fig.1e,f,g and SI Fig.1b,c; Movie S3). This confirms that RhoA affects the magnitude but not the handedness of chiral flows.

### Chiral acto-myosin flows depend linearly on cortical Lifeact::mKate2 intensity

Since the presence of high amounts of Lifeact::mKate2 reverses the handedness of chiral flows, we asked if there is a specific dosage of Lifeact::mKate2 at which the reversal takes place. To evaluate this, we compared chiral velocities 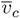 obtained in three transgenic worm strains [12, 34] that express varying amounts of the identical Lifeact::mKate2 construct (SI Fig.1d; also see Methods; Movies S1, S4). We find that the chiral velocity 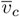 scales linearly with the amount of cortical Lifeact:mKate2 as determined by fluorescence intensity measurements (Pearson’s Correlation Coefficient *R* = 0.64, *p* − *value* << 0.001, Fig.1h; see Methods). We note that chiral reversal of cortical flows, indicated by a change of sign of 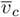 from negative to positive, occurs at a cortical Lifeact::mKate2 intensity of 0.5 × 10^8^ *A*.*U*., which is ∼50% of the average cortical intensity in Lifeact++ zygotes (1.12× 10^8^ *A*.*U*., Fig.1h). Chirality reversal is unlikely to result from an off-target effect due to genomic integration of Lifeact, given that the three strains used here were generated using different integration methods at different genomic locations ([12, 34]; see Methods). Taken together, these results are consistent with the interpretation that the magnitude of active torque depends on the amount of cortical Lifeact::mKate2, with the handedness of active torques reversing at a specific dosage.

**Fig. 1:**
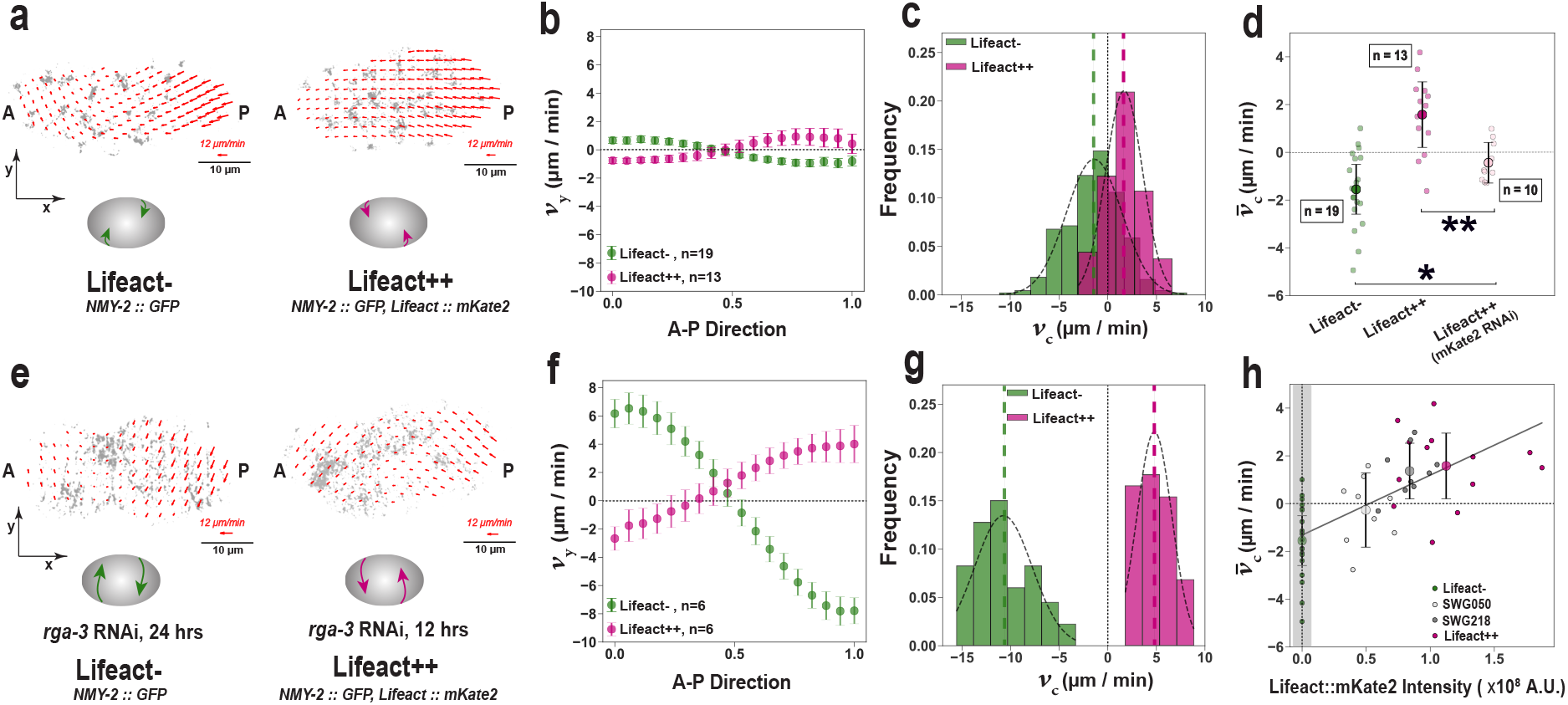
High expression of Lifeact::mKate2 reverses handedness of cortical flows during polarity onset in C. elegans zygotes. Representative fluorescence microscopy images of cortical non-muscle myosin 2 (NMY-2::GFP) in zygotes of Lifeact- and Lifeact++ strains during onset of polarity. The long axis (A-P direction) of the embryo forms the x-axis. Red arrows indicate velocity vectors from PIV. (b) The average flow component along the y-axis (v_y_), plotted against relative position along the A-P direction of the zygote. The v_y_ profiles were averaged over 19 Lifeact- and 13 Lifeact++ zygotes respectively (see Methods). (c) Instantaneous chiral velocity v_c_ distributions in Lifeact- and Lifeact++ zygotes. v_c_ was measured for 542 frames across 19 Lifeact- and 306 frames across 13 Lifeact++ zygotes. In (c) and (g) the black dashed curves represent gaussian fits for the respective distributions. The mean with 99% confidence interval from the gaussian fit is -1.39 ± 0.32 µm/min for Lifeact- and +1.68 ± 0.28 µm/min for Lifeact++ respectively. (d) Rescue of cortical flow handedness, indicated by negative chiral velocity 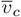, was achieved in 7/10 Lifeact++ zygotes upon mKate2(RNAi). A one-sided Mann– Whitney U Test was used for assessing significance. * refers to p < 0.05; ** refers to p < 0.01. (e) Fluorescence images of NMY-2::GFP overlaid with PIV flow profiles during polarity onset under rga-3(RNAi) . Duration of RNAi was 24 hrs for Lifeact- and 12 hrs for Lifeact++ zygotes. (f) Average v_y_ of the flow fields under rga-3(RNAi) plotted against the relative position along the A-P direction. v_y_ profiles were averaged over six Lifeact- and six Lifeact++ zygotes respectively (see Methods). (g) v_c_ distributions under rga-3(RNAi). 76 frames across six Lifeact-, and 99 frames across six Lifeact++ zygotes were analysed. The mean with 99% confidence interval from the gaussian fit is -10.75 ± 0.90 µm/min for Lifeact-zygotes and +4.80 ± 0.48 µm/min for Lifeact++ zygotes respectively under rga-3(RNAi). (h) Magnitude of 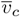 during polarity onset linearly depends on the intensity of cortical Lifeact::mKate2 in the zygote. Scatter plot shows measured 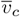 and cortical Lifeact::mKate2 intensity for each individual zygote from 4 transgenic C. elegans strains : Lifeact-, SWG050, SWG218 and Lifeact++ (see Methods). The last three strains express different levels of the identical Lifeact::mKate2 construct (see SI Fig.1d). Intensity of Lifeact::mKate2 in Lifeact-zygotes was assumed 0 (solid gray bar) and included to obtain the linear fit. Error bars refer to the SEM in (b,f) and 99% confidence interval to the mean in (d,h).

**SI Fig. 1:**
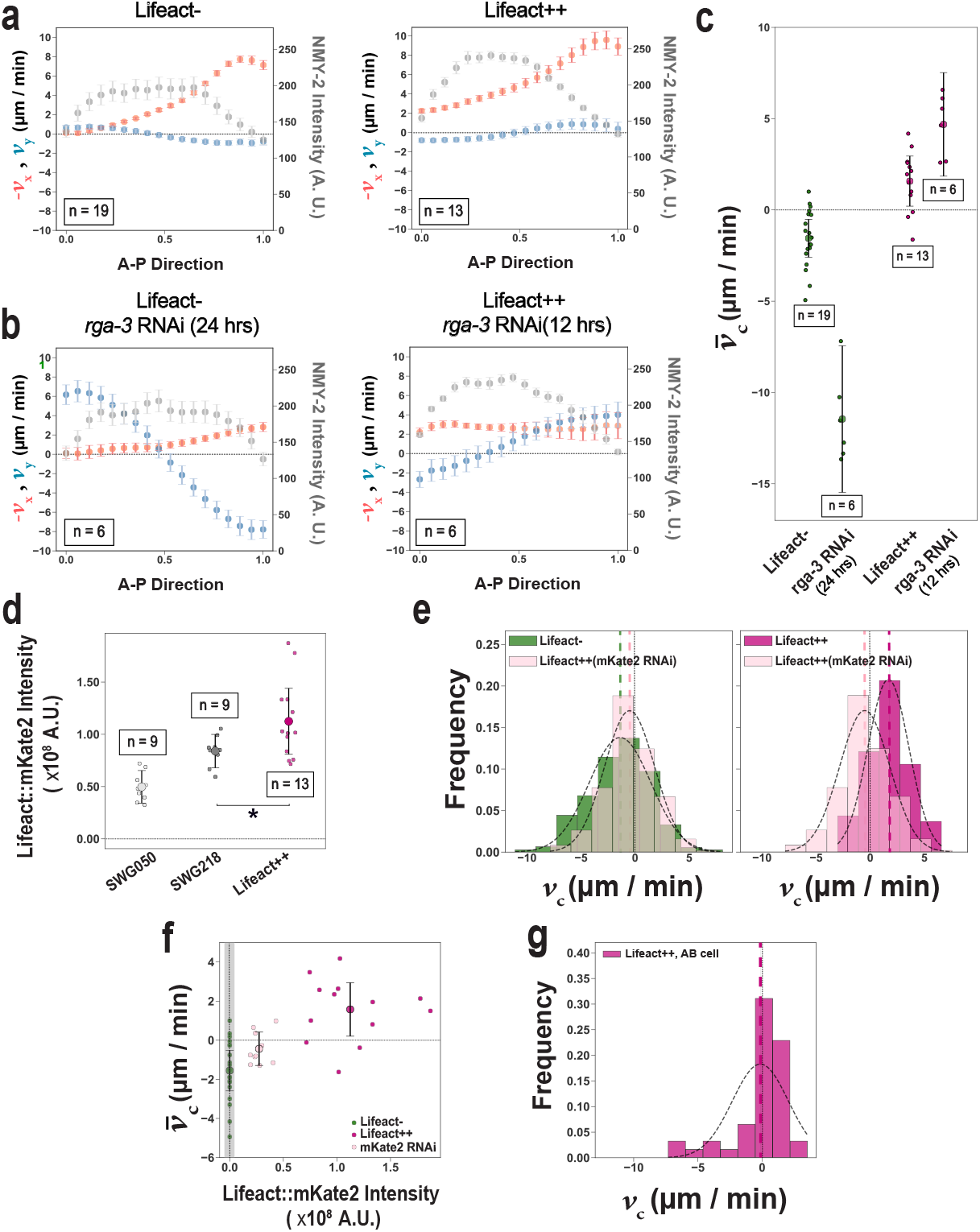
Supplementary Information on cortical flows during polarity onset in C. elegans zygotes. (a) Dual y-axis plots showing cortical velocity and myosin intensity profiles in zygotes. The left y-axis represents the average cortical flow component along the A-P direction (v_x_) and orthogonal to the A-P direction (v_y_), while the right y-axis shows cortical NMY-2::GFP intensity; all plotted against relative position along the A-P direction of the zygote represented along the x-axis of the plot. Cortical flow and myosin intensity profiles were averaged over 19 Lifeact- and 13 Lifeact++ zygotes (see Methods). Error bars denote the SEM. (b) The v_x_, v_y_ and NMY-2::GFP intensity profiles during polarity onset under rga-3(RNAi) condition. All profiles were averaged over six Lifeact- and six Lifeact++ zygotes. (c) Chiral velocity 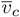 measured for Lifeact- and Lifeact++ zygotes, for control and rga-3(RNAi) condition. Error bars denote 99% confidence to the mean. (d) Cortical Lifeact::mKate2 intensity measured in three transgenic worm strains expressing the identical Lifeact::mKate2 construct. In order from left to right; SWG050 (n=9), SWG218 (n=9) and Lifeact++ (n=13) zygotes (for more details, see Methods). Error bars denote 99% confidence to the mean. A one-sided Mann–Whitney U Test was used to assess whether SWG218 had lower cortical Lifeact::mKate2 intensity compared with Lifeact++ (p = 0.041; indicated by *). (e) Right panel; distributions of v_c_ for Lifeact++ zygotes under mKate2(RNAi) condition and Lifeact-zygotes. The black dotted curves denote the gaussian fits for the respective distributions. Left panel; likewise for Lifeact++ zygotes, control and under mKate2(RNAi) condition. The mean and 99% confidence interval from the gaussian fit is -0.51 ± 2.31 µm/min for Lifeact++ under mKate2(RNAi). (f) Chiral velocity 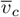 plotted against cortical Lifeact::mKate2 intensity for Lifeact-, Lifeact++ and Lifeact++ zygotes under mKate2(RNAi). (g) Distribution of v_c_ for the AB cell in the Lifeact++ strain. The black dotted curve denotes the gaussian fit for the distribution. The mean and 99% confidence to the mean is -0.14 ± 0.89 µm/min.

### Lifeact reverses the handedness of acto-myosin flows during LR symmetry breaking

Chiral cortical flows have been implicated in LR symmetry breaking in the nematode embryo by a skew of the division axes at the 4-6 cell stage (Fig.2a) [2]. Given that high amounts of Lifeact::mKate2 reverse the handedness of chiral flows in the *C. elegans* zygote, we asked if a similar reversal occurs during LR symmetry breaking at the 4-6 cell stage. We speculated that cortical flows with reversed handedness during ABa and ABp cell divisions would skew their division axes in the opposite direction, thereby establishing reversed LR asymmetry. To this end, we analysed cortical flow fields during division of the ABa cell (see Movies S5, S6), and measured instantaneous chiral velocities *v*_*c*_ in Lifeact- and Lifeact++ embryos. *v*_*c*_ for the ABa cell is defined as 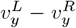, where 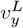 and 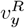 denote the *y*-component of the space-averaged instantaneous velocities in the left (L) and right (R) domains ([2, 16]; see Methods). We find that *v*_*c*_ is distributed around 0 for the Lifeact++ condition (−0.05 ± 0.14 *µm/min*; mean with 99% confidence from the gaussian fit; Fig.2d) with an increase in positive values compared with the Lifeact-condition (−2.91 ± 0.50 *µm/min*, Fig.2d, SI Fig.2b). Note that an abundance of positive *v*_*c*_ under the Lifeact++ condition indicates a high frequency of instantaneous reversals in flow handedness during division of the ABa cell. Measurement of chiral velocity 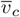, by averaging *v*_*c*_ over time for each embryo (for more details, see Methods) reveals, that indeed, 21/38 (∼ 55%) embryos with the Lifeact++ condition display positive 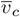, in contrast to all Lifeact-embryos that display negative 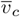 (Fig.2e) [2, 16]. From this, we conclude that high amounts of Lifeact::mKate2 in the cortex can reverse cortical flow handedness during division of the ABa cell.

The ABp cell divides simultaneously as the ABa cell (Fig.2a; Movies S7, S8). We find that 13/38 (∼ 34%) embryos of Lifeact++ display positive 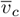 during the ABp cell division, compared with negative 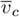 for all the Lifeact-embryos (Fig.2i, SI Fig.2c). Taken together with our previous findings, we conclude that the reversal in flow handedness observed in *C. elegans* zygotes with high amounts of cortical Lifeact::mKate2 holds true at the 4-6 cell stage during LR symmetry breaking. Note that a similar effect can be observed in the AB cell during its division at the 2-3 cell stage ([33]; see SI Fig. 1g).

**Fig. 2:**
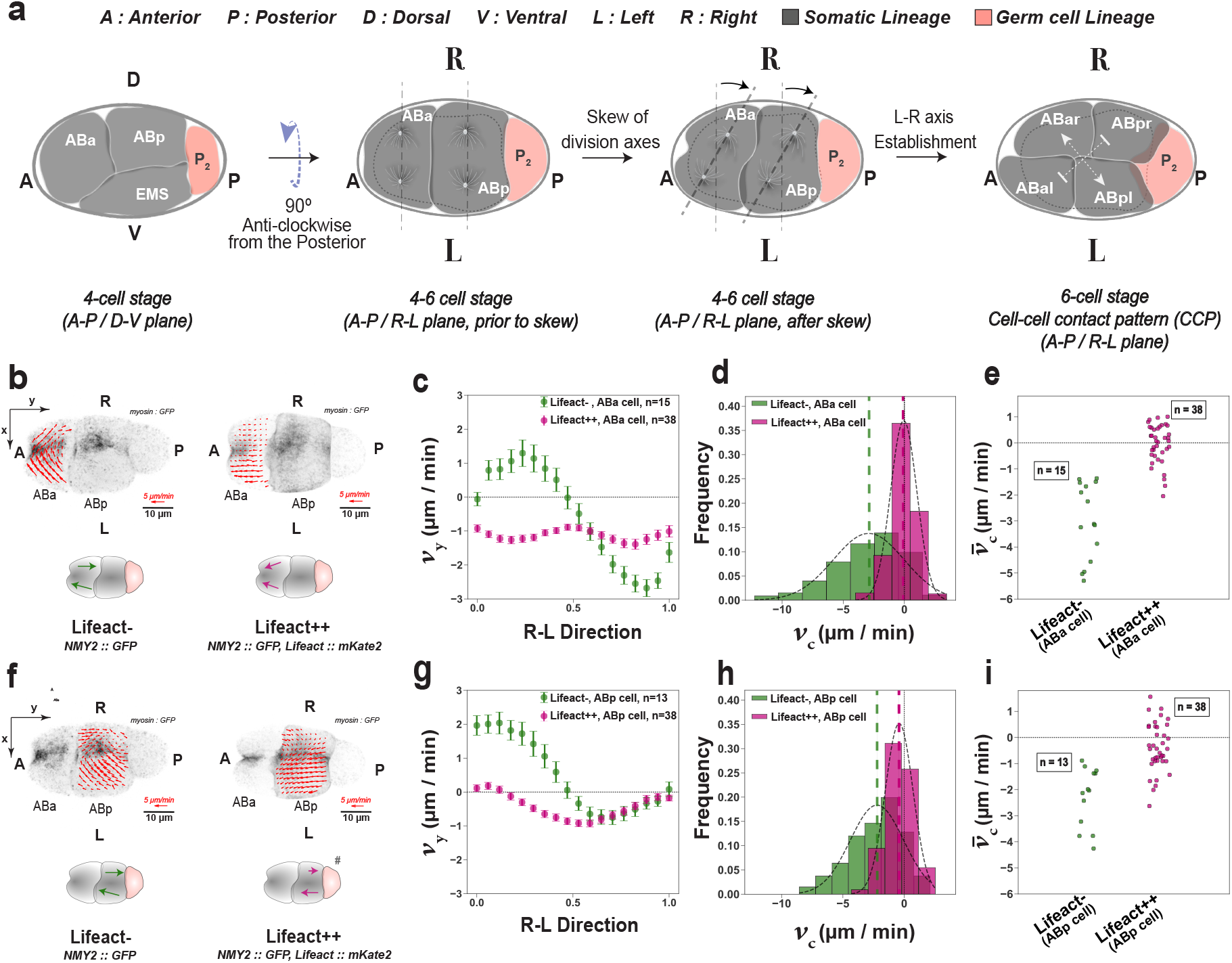
High amounts of Lifeact::mKate2 reverses the handedness of cortical flows during LR symmetry breaking at the 4-6 cell stage. (a) Schematic showing the 4-6 cell stage transition in C. elegans embryos. The resulting cell-cell contact pattern (CCP) at the 6-cell stage is marked by the dashed arrows, viewed from the dorsal side in the A-P/R-L plane. Five cells, namely ABar and ABal (the ABa daughters), ABpr and ABpl (the ABp daughters) and P_2_ can be observed in the dorsal view. The sixth cell, EMS, is on the ventral side and is marked by the gray dashed cell behind the viewing plane. Fluorescent images of NMY-2::GFP overlaid with flow profiles (red arrows) during ABa division for representative embryos of Lifeact- and Lifeact++. The R-L direction of the embryo serves as the x-axis, while the y-axis is along the A-P direction. (c) The average v_y_ component of the cortical flow during the ABa division plotted against relative position along the R-L direction of the ABa cell. The profiles represent averages respectively over 15 Lifeact- and 38 Lifeact++ embryos. (d) Histograms of instantaneous chiral velocity v_c_ for Lifeact- and Lifeact++ embryos during ABa division. v_c_ was measured respectively for 231 frames across 15 Lifeact-, and 391 frames across 38 Lifeact++ embryos. 231/391 frames showed positive v_c_ for Lifeact++ compared with 41/231 frames for Lifeact-. Black dashed curves denote gaussian fits on the respective distributions. The mean with 99% confidence from the gaussian fits are -2.91 ± 0.50 µm/min for Lifeact- and -0.05 ± 0.14 µm/min for Lifeact++. (e) Chiral velocity 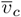, averaged over time for each embryo of Lifeact-(n=15) and Lifeact++ (n=38) during ABa division (see Methods). (f,g,h,i) show the equivalent for the ABp cell division, as (b,c,d,e) do for the ABa cell, in Lifeact-(n=13) and Lifeact++ embryos (n=38). v_c_ during ABp division was measured for 199 frames across 13 Lifeact-embryos, and 384 frames across 38 Lifeact++ embryos. 148/384 frames showed positive v_c_ in Lifeact++ compared with 28/199 frames in Lifeact-during ABp division. The mean with 99% confidence from the gaussian fit is -2.22 ± 0.40 µm/min for Lifeact- and -0.41 ± 0.15 µm/min for Lifeact++ respectively. The schematics in (b,f) show direction of cortical flows in the ABa and ABp cells. Lifeact++ embryos displayed inconsistency in cortical flow directionality during ABp division (see SI Fig.2a; Movie S8). The respective schematic indicated by # only shows the flow directional of the representative embryo above.

**SI Fig. 2:**
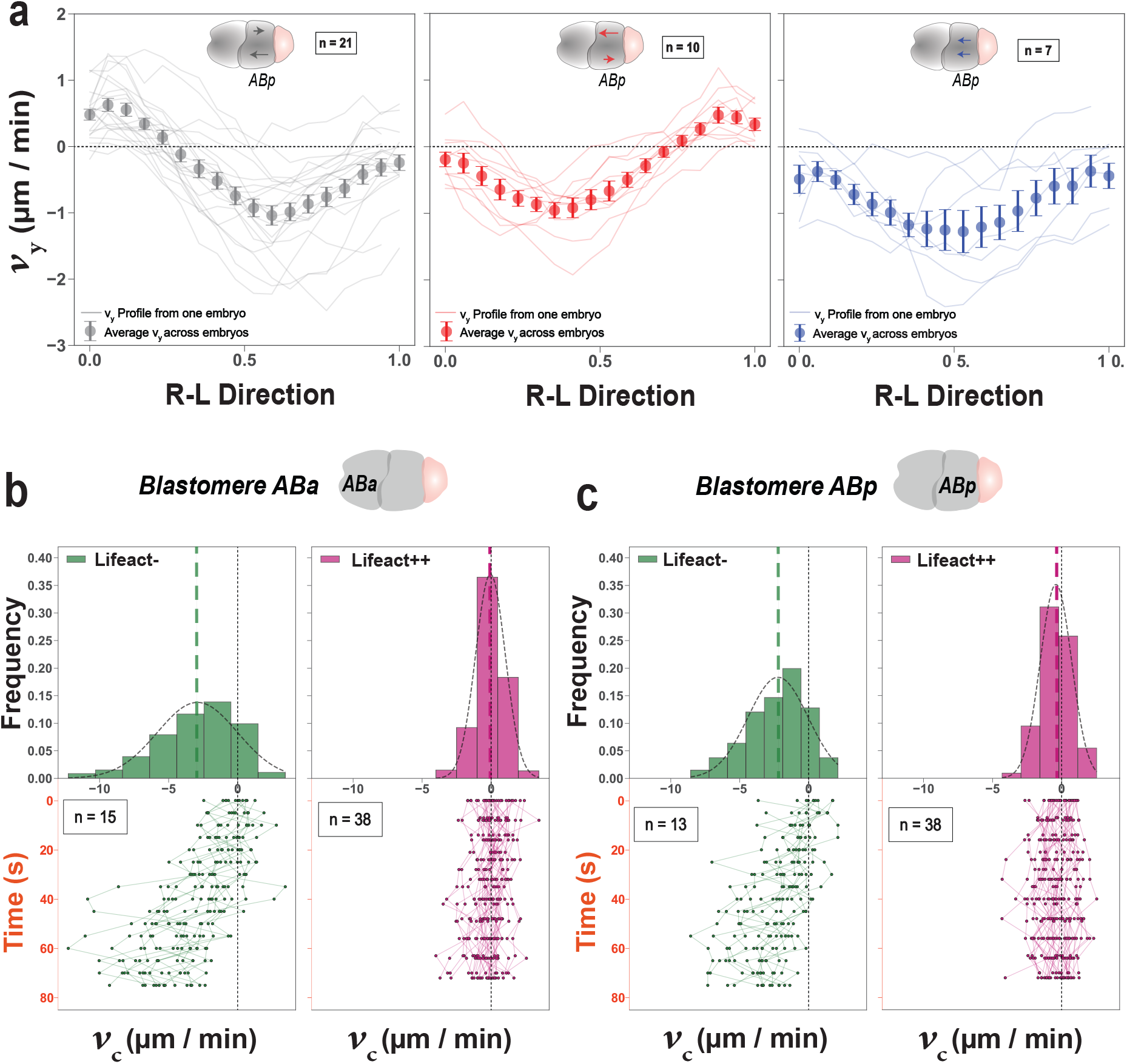
Supplementary information on ABa and ABp flow profiles during cytokinesis. (a) The average v_y_ component of the cortical flow during ABp division for each individual embryo is plotted against relative position along the R-L direction of Lifeact++(n=38). Left panel; 21/38 embryos have cortical counter-rotation flows between left and right halves of the ABp with the stereotypic directionality. Central panel; 10/38 embryos show reversed directionality of rotational flows between the left and right halves of the ABp cell cortex. Right panel; 7/38 embryos show anterior-ward flows on both halves of the cortex during ABp division. Error bars denote SEM. (b) Data used for v_c_ distributions in Fig.2e. Histograms contain time evolution of v_c_ for each embryo during ABa division for Lifeact-(left-panel; 231 frames across 15 embryos) and Lifeact++ (right-panel; 391 frames across 38 embryos). (c) Likewise as (b) for the ABp division, corresponding to the respective histograms in Fig.2i (left panel, 199 frames across 13 Lifeact-embryos; right panel, 384 frames across 38 Lifeact++ embryos).

### Cortical flow handedness is instructive for the 6-cell stage contact pattern

The ABa and ABp division skew results in asymmetric positioning of their daughter cells within the embryo [1]. The left daughter cells of the ABa and ABp (ABal, ABpl) are more anteriorly positioned compared to their right counterparts (ABar, ABpr; see Fig.2a) [1]. Together, this gives rise to a specific cell-cell contact pattern (CCP) at the 6-cell stage of the embryo [1], where ABar and ABpl are in contact with each other (Fig.2a, right-most panel; pointed arrow heads), while ABal and ABpr are not (Fig.2a, flat arrow heads). This LR asymmetric CCP is stereotypic in wild-type worms, and is referred to as the dextral CCP [1, 21, 40]. In 1991, W. B. Wood generated the opposite (sinistral) contact-pattern at the 6-cell stage, by pushing the ABa cell with a micro needle during its division and reversing the division skew. Here, we asked whether cortical flows with reversed handedness 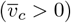 during the ABa division generate a sinistral CCP. To this end, we divided Lifeact++ embryos according to the sign of their 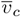 during ABa division, and examined the CCP that resulted in each embryo at the 6-cell stage. We find the probability of sinistral CCP with 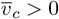 is 0.63 (0.38,0.84) (central estimate with 95% exact binomial confidence intervals within brackets; for more details, see Methods), in contrast to 0.12 (0.01,0.36) for embryos with 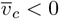 (SI Fig.3a, left panel). Likewise, dividing the embryos by sign of 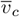 during the ABp division, we find that sinistral CCP has an increased probability of 0.73 (0.39,0.94) for 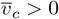, compared with 0.20 (0.07,0.41) for 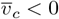 (SI Fig.3a, right panel). We next divided our observed range of 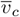 into 0.5 *µm/min* bins, and determined in each bin, the fraction of embryos with dextral and sinistral CCP. We find that the probability of sinistral CCP becomes non-zero with increasing 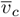, showing significant increase when 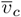 changes to positive values. This demonstrates a gradual shift in the 6-cell CCP preference as the flow handedness reverses (Fig.3b; lower panel). Accordingly, sinistral 6-cell embryos showed reversed anti-symmetry in *v*_*y*_ along the R-L direction (Figs. 3b,c) and negative chirality index *c* during division of the ABa (*c* − = 0.41(− 0.69, − 0.16) for sinistral CCP; 0.30(0.07, 0.70) for dextral CCP) and ABp cells (*c* − = 0.30(− 1.23, − 0.07) for sinistral CCP; 2.72(1.28, 5.14) for dextral CCP). Consistent with these results, embryos with the sinistral CCP displayed a positive median 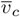 of 0.26 (0.01,0.65) *µm/min* (25th percentile, 75th percentile within brackets) during the ABa cell division, and 0.42 (-0.30,0.71) *µm/min* during the ABp cell division (Fig.3b, upper panel). Furthermore, embryos with the dextral CCP displayed a negative median 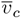 of -0.27 (-0.69,0.22) *µm/min* during the ABa cell division, and -0.84 (-1.22,-0.52) *µm/min* during the ABp division. Taken together, our results are consistent with the mechanism that the handedness of cortical flows during ABa and ABp divisions instructs the handedness of the CCP at the 6-cell stage.

**Fig. 3:**
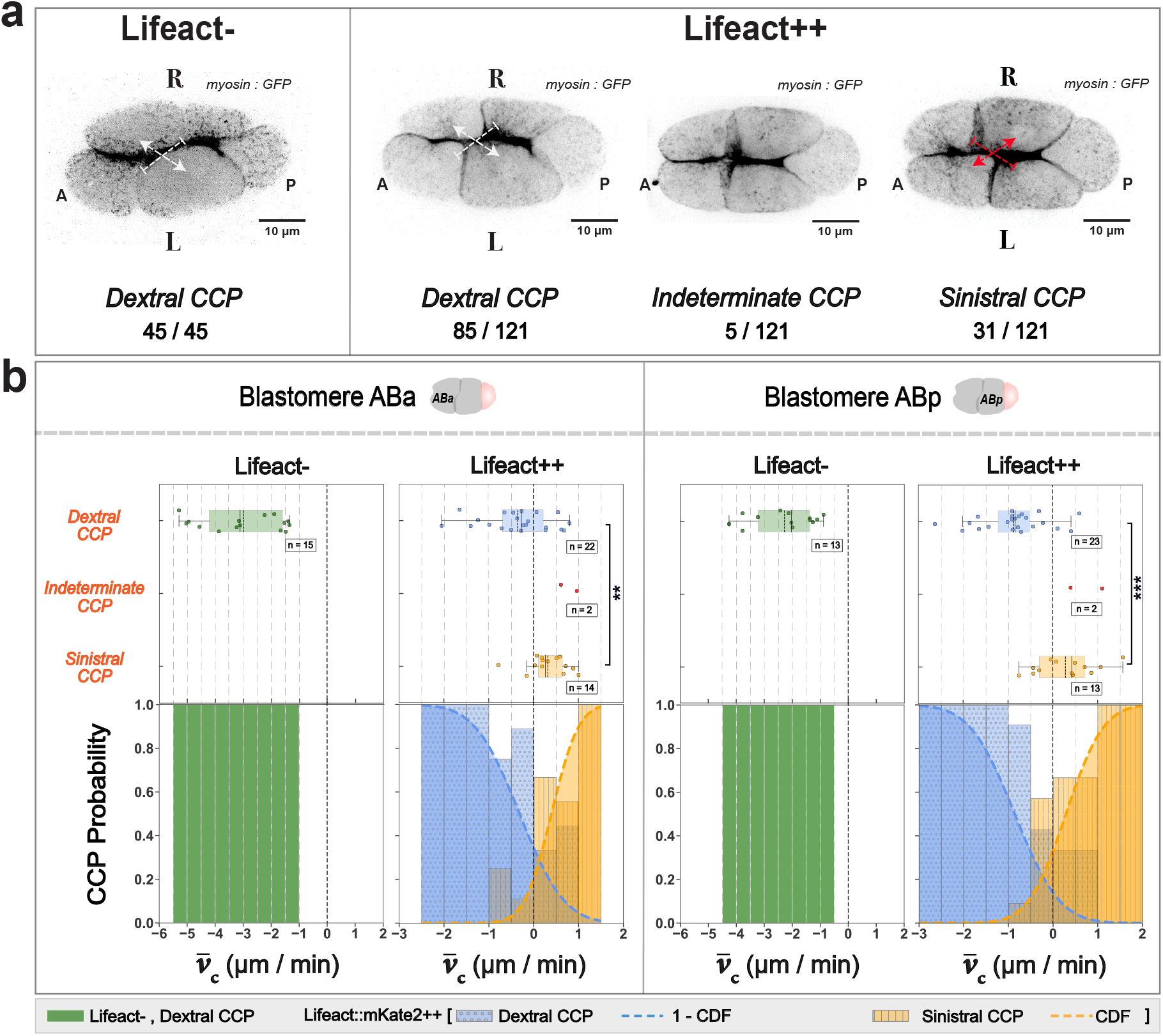
Reversed handedness of cortical flows during 4-6 cell transition can reverse the handedness of the 6-cell stage CCP. (a) Effect of excess Lifeact::mKate2 on the handedness of the 6-cell stage CCP in C. elegans embryos. Left panel shows the stereotypic dextral CCP at the 6-cell stage of Lifeact-embryos (45/45). Right panel shows Lifeact++ embryos with the three categories of CCP observed at the 6-cell stage and their respective sample sizes. White arrows denote the dextral CCP; red arrows denote the sinistral CCP. (b) Likelihood of sinistral 6-cell stage CCP increases with increasingly reversed chiral velocity 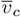 during ABa and ABp divisions. Upper left panel; 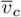 plotted on the x-axis and classified according to the 6-cell stage CCP. Lower left panel; 6-cell stage CCP probability plotted for each 0.5 µm/min bin along the observed range of 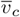 during the ABa cell division. Right panels as left panels but for the ABp cell division. ** denotes p < 0.01; *** denotes p < 0.001. A two-sided Mann–Whitney U test was performed to test for significance. Shaded boxes in the whisker plots denote the range of the data between the 25th and the 75th percentiles. Black solid and dashed lines inside the shaded boxes denote the median and the mean respectively. The gray dashed lines denote the 0.5 µm/min 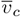 bins.

Following the establishment of the 6-cell stage CCP, LR asymmetric patterning occurs and distinct cell fates emerge on the left and right side of the developing embryo [1, 21–25]. This led us to enquire the frequency of distinct CCPs in the Lifeact++ and Lifeact-populations. To this end, we investigated 6-cell stage CCP and classified them into dextral, sinistral, or indeterminate (all four daughter cells are in contact); see Fig.3a; Movie S9). We find that ∼70% (85/121) of the Lifeact++ embryos displayed dextral CCP, ∼ 25% (31/121) displayed sinistral CCP, and ∼5%(5/121) displayed indeterminate CCP at the 6-cell stage, compared with 100% (45/45) dextral CCP in the Lifeact-embryos (Fig.3a). Together with our previous findings, we conclude that Lifeact::mKate2 not only affects the handedness of cortical flows, but also significantly increases the frequency of the sinistral CCP at the 6-cell stage.

**SI Fig. 3:**
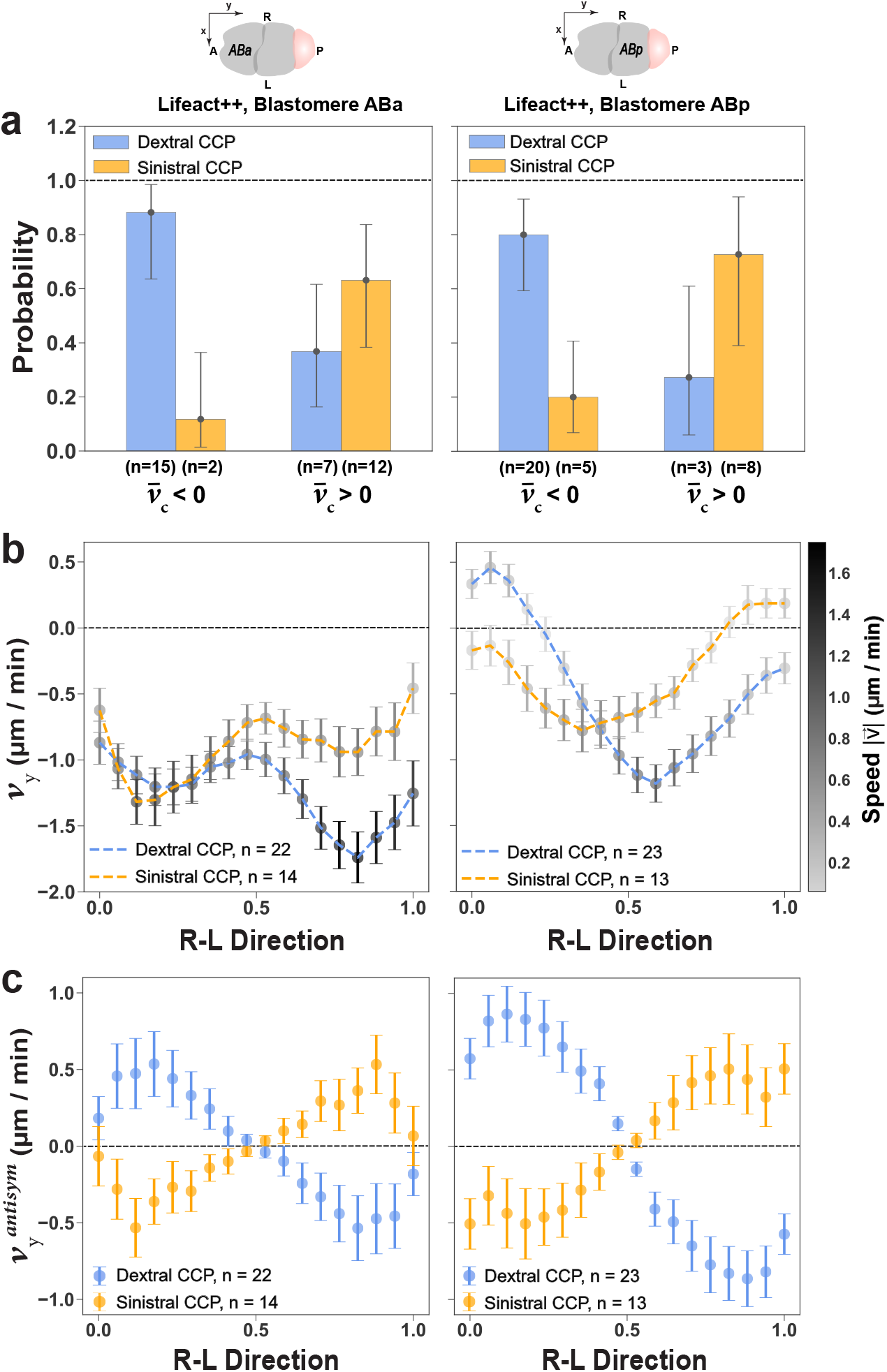
(a) Probability of dextral or the sinistral CCP at the 6-cell stage for a given handedness of cortical flows at the 4-6 cell stage. Embryos classified by the sign of chiral velocity 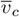 during division of the ABa cell (left panel), and the ABp cell (right panel). Error bars denote Clopper–Pearson exact 95% confidence intervals (see Methods). (b) The average y-component of cortical flows v_y_ plotted along the R-L direction of the ABa cell (left panel), and the ABp cell (right panel) for embryos with dextral and sinistral CCPs at the 6-cell stage. Color bar on the right denotes speed of cortical flows for each respective group along the R-L direction. (c) The average anti-symmetric y-component of the cortical flows 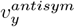 plotted along the R-L direction of the ABa cell (left panel), and the ABp cell (right panel) for embryos with dextral and sinistral CCP at the 6-cell stage. For details on calculation of 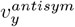, see Methods. Error bars in (b) and (c) denote the SEM. Note that the embryo ensembles used in (a),(b) and (c) do not include those with indeterminate CCP at the 6-cell stage.

### Reversal of cortical flow handedness results in *situs inversus* adults

The handedness of the 6-cell CCP determines the LR axis, and embryos with a sinistral CCP have been reported to develop into *situs inversus* adults [1, 21, 40]. Therefore, we expect embryos with reversed cortical flows and a sinistral CCP to develop into *situs inversus* adults. To evaluate this, we isolated Lifeact++ embryos that had the dextral or sinistral CCP, and subsequently determined the LR body axis of the resulting adult worm, by examining the relative position of its anterior gonad and intestine (see Methods) [1, 21]. We find that 11/11 embryos with sinistral CCP developed into *situs inversus* adults, while 13/13 embryos with dextral CCP displayed the stereotypical body plan (Fig.4a, see also SI Fig.4; ; Movie S10). Note that the brood size of *situs inversus* hermaphrodites does not significantly differ from their unreversed counterparts and agrees with [1] (Fig.4b). Furthermore, the fraction of dextral vs sinistral worms does not differ significantly in the brood of dextral/sinistral worms [40] and compares to ∼3% of *situs inversus* worms (32 in 885) in a regular culture of Lifeact++ worms. In conclusion, we show that reversing the handedness of cortical flows during embryonic LR symmetry breaking reverses body axis handedness, generating *situs inversus* worms.

**SI Fig. 4:**
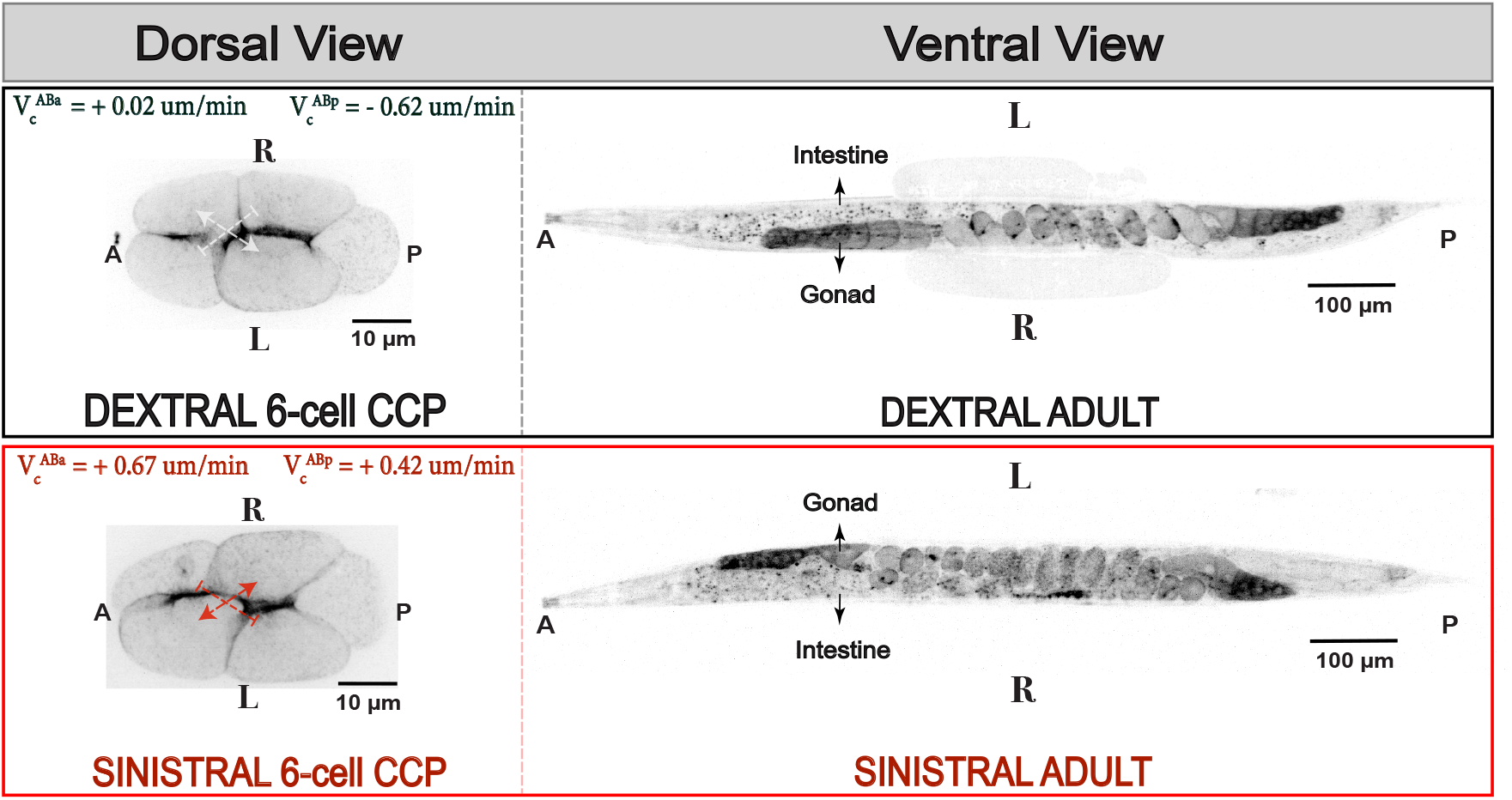
LR handedness of C. elegans hermaphrodites, as viewed from the ventral side. Upper panel; a 6-cell stage embryo mounted on the dorsal side showing dextral CCP. The reported values are chiral velocities for cortical flows during ABa and ABp cell divisions in the representative embryo. The corresponding adult showing the ventral gonad and intestine on the right and the left sides respectively, when viewed from the ventral side. Lower panel; likewise for a 6-cell embryo with sinistral CCP and the corresponding adult showing sinistral LR handedness in its gonad-gut placement on the ventral side.

## Discussion

The handedness of the *C. elegans* body plan in wild type population is essentially invariant, with approximately less than 1 in 15,000 worms being *situs inversus* [21]. Previous studies have identified factors such as mechanical perturbation [1], cold shock [40], removal of the egg shell [40] and randomization of early spindle orientations [21] as means to generate *situs inversus* worms. Notably, Bergmann et.al.’s study [21] showed that randomization of the ABa and ABp spindle orientations randomizes the handedness of the LR axis. Work done by Naganthan et.al. [2] in 2014 identified chiral flows in the cortex of the ABa cell as the upstream driver of the relevant spindle skew during its division. This study now establishes active torques generated in the actomyosin cortical layer as instructive for LR axis specification. Reversal of active torque handedness can reverse the 6-cell stage CCP, giving rise to ∼ 1 in 30 *situs inversus* worms. We show that high amounts of Lifeact::mKate2 in the cortex drive this reversal of active torque handedness. Since the acto-myosin cytoskeleton can participates in chiral symmetry breaking in vertebrates [17, 41] and invertebrates [2, 5, 7, 8, 11, 16], it will be interesting to see if using Lifeact::mKate2 can reverse body axis handedness in these systems as well.

It will also be interesting to shed light onto the molecular mechanism by which Lifeact::mKate2 reverses the handedness of cortical active torques. Lifeact’s binding site on F-actin is same as that of myosin and cofilin [28], both of which have been implicated in active torque generation [2, 7, 8, 12, 17, 42, 43]. Non-muscle myosin II drives active torques by cross-linking anti-parallel actin filaments [2, 19], as well as in concert with the actin elongator formin [12]. Cofilin, on the other hand, severs F-actin [44] and changes the helical pitch of actin filaments upon binding [42, 43]. Studies using only Lifeact as well as Lifeact constructs fused with different fluorophores have shown that filament severing and cofilin binding to actin decrease at high concentrations *in-vitro* [26, 28]. Lifeact also affects actin assembly and elongation [26]. Our work now provides evidence that in addition to changing essential properties of the cytoskeleton such as its turnover and contractility, Lifeact can also reverse the handedness of active torques and flows in the cortex. Future work will elucidate the molecular mechanism by which Lifeact changes the handedness of active torque generation, and shed light onto how chirality at the mesoscale emerges from molecular interactions.

## Methods

### Worm Strains

Transgenic worm strains used in this study: LP133 (*nmy-2(cp8[nmy-2::gfp +LoxP])I; unc119(ed3)III*)[2][45][16][12], SWG050 (*gesIs003 [Pmex-5::Lifeact::mKate2::nmy-2UTR, unc-119+]*)[12], SWG218 (*nmy-2(cp13[nmy-2::GFP + LoxP]) I; gesSi089[Pmex-5::Lifeact::mKate2; unc 119 (+)] II; unc-119(ed3) III*) and SWG007 (*nmy-2(cp8 [nmy-2::GFP unc-119+]) I; gesIs001 [Pmex-5::Lifeact::mKate2::nmy-2UTR, unc-119+]*)[34]. LP133 is referred to as Lifeact-in this study. SWG050, SWG218 and SWG007 express the identical Lifeact::mKate2 construct with a 66 bp linker between the actin probe and the fluorophore. SWG001 (*gesIs001[Pmex-5::Lifeact::mKate2::nmy-2UTR, unc-119+]*) [34] was crossed with LP133 to obtain SWG007. Lifeact++ refers to SWG007 in this study. SWG050 and SWG001 were obtained by bombardment with Lifeact::mKate2 construct (*MosSCI-mex-5pCter-Lifeact-mKate2*). SWG218 was obtained by introducing the Lifeact::mKate2 construct as a MosSCI insertion in SWG213 (*nmy-2(cp13[nmy-2::GFP + LoxP]) I; ttTi5605 II; unc-119(ed3) III*). All strains were cultivated at 20^*°*^C on OP50 plates, as indicated in Brenner 1974 [46].

### RNA Interference

RNA interference (RNAi) experiments were performed by feeding the worms, according to Timmons et al 2001 [47]. Worms were incubated on plates with NGM agar, 1 mM isopropyl-*β*-D-thiogalactoside (IPTG) and 50 *µg/ml* ampicillin. RNAi feeding time was defined between transfer of worms to the RNAi feeding plates and the putative time of imaging the embryos after dissection. The embryos were imaged directly after dissection, leaving a gap of maximum 5 minutes between picking the worms from the RNAi feeding plates into M9 buffer and imaging. The *mKate2(RNAi)* feeding clone was obtained from the Ahringer Library [48, 49].

### Imaging of *C. elegans* embryos

L4 larvae were picked 24 hrs before imaging. For imaging *C. elegans* zygotes and 2-3 cell embryos, adults were dissected in a mixture of M9 buffer and 20 *µm* polystyrene beads (Polysciences), mounted onto a coverslip and sealed with wax. Image acquisition was carried out using a Yokogawa CSU-X1 scan head with a Hamamatsu ORCA-Flash 4.0 camera, in combination with two objectives, namely, the Zeiss C-Apochromat 100x/1.42 NA(oil) objective, and the Zeiss C-Apochromat 63x/1.2 NA(water) objective. Majority of the zygotes and the 2-3 cell stage embryos were imaged with the 100x/1.42 NA(oil) objective, while the 4-6 cell stage embryos were imaged using the 63x/1.2 NA(water) objective. Three z-slices 0.5 *µm* apart, starting from the cortex inwards, were imaged every 5 seconds to image cortical flows in the zygotes and the AB cell at the 2-3 cell stage. For imaging *C. elegans* embryos at the 4-6 cell stage, a different mounting technique was used. The adults were dissected in M9 buffer, and 4-cell embryos were identified. Prior to imaging, the 4-cell embryos were turned to their dorsal side using an eye-lash tool. The samples were not sealed with a cover-slip to maintain the required orientation. Note that Lifeact++ embryos were imaged for body plan handedness after they had been in culture for 8 weeks, as an increase in the frequency of reversals was noted over time. Initially during the study, cortical flows were imaged using a z-stack comprising of 23 z-slices 0.5 *µm* apart every 5 seconds in Lifeact-embryos, and every 7 seconds in Lifeact++ embryos. However, embryos often exhibited a rotation along the A-P axis during late 4-6 cell stage, forcing parts of the ABa and ABp cells out of the imaging plane during establishment of the 6-cell stage contact pattern. To account for this, we imaged a z-stack comprising of 29 z-slices 0.75 *µm* apart every 5 seconds in Lifeact-embryos, and every 8 seconds in the Lifeact++ embryos. Note that the shorter z-stack is sufficient for imaging cortical flows but compromises the quality of imaging at the 6-cell stage. Since analysis of cortical flows was carried out during early cytokinesis at the 4-6 cell stage, images obtained from both the aforementioned settings were included for analysis. For Particle Image Velocimetry (PIV), maximum intensity projections were obtained over a distance of 11-15 *µm* starting at the cortex on the dorsal side (all 23 z-slices 0.5 *µm* apart; first 15-20 z-slices 0.75 *µm* apart).

### Imaging of Adults for Gut-Gonad chirality

After imaging ABa and ABp divisions during the 4-6 cell stage, each 6-cell embryo was pipetted out of the buffer using a manual mouth-pipette onto a fresh OP50 plate, and stored at 20^*°*^C until it hatched and developed into an adult hermaphrodite. For imaging the gut-gonad twist in the adult body plan, the hermaphrodite was transferred from the OP50 plate onto an agar pad on a glass slide and paralyzed in 15 *µl* of 1 mM Levamisole (Sigma Aldrich; CAS No. 16595-80-5) for 15 minutes. Upon immobilization, the worm was turned to have a dorsal-side up or a ventral-side up orientation using an eye-lash tool to ensure observation of the gonad and the gut simultaneously. A cover slip was placed on the sample and imaged using a Zeiss Plan-Apochromat 10x/0.45 objective. A z-stack comprising of 52 z-slices 1*µm* apart was acquired. A ventral-side up view of the worm can be obtained by turning the worm such that its uterus faces the cover slip [1][40][21]. The dorsal-side up view, therefore, can be obtained by facing the worm uterus directly away from cover slip. In the ventral-side up view, the gut-gonad placement in the region between the pharynx and the uterus of the worm was checked. For a dextral adult turned on its ventral side, the anterior-proximal gonad with the oocyte train must be located on the worm’s right side, while the intestine must be on the worm’s left side. If the arrangement was flipped, the worms were classified as *situs inversus* or sinistral adult, as illustrated by the maximum intensity projections shown in Fig.4 [1][40][21]. In the dorsal-side up view, the gut-gonad arrangement flips, as the worm is turned 180^*°*^ from its ventral view (Fig.4a shows maximum intensity projections). As the embryo was turned on its dorsal side for observation of LR symmetry breaking, we included the dorsal view of the worms for a direct comparison (Fig. 4a). Note that if the worm rolled over to its left or right side upon placement of the cover slip, the gut-gonad chirality could still be analysed with the aid of the 51 *µm* wide z-stack of the adult. In such situations, the dorso-ventral axis was assigned by the placement of the the distal gonad and the uterus.

**Fig. 4:**
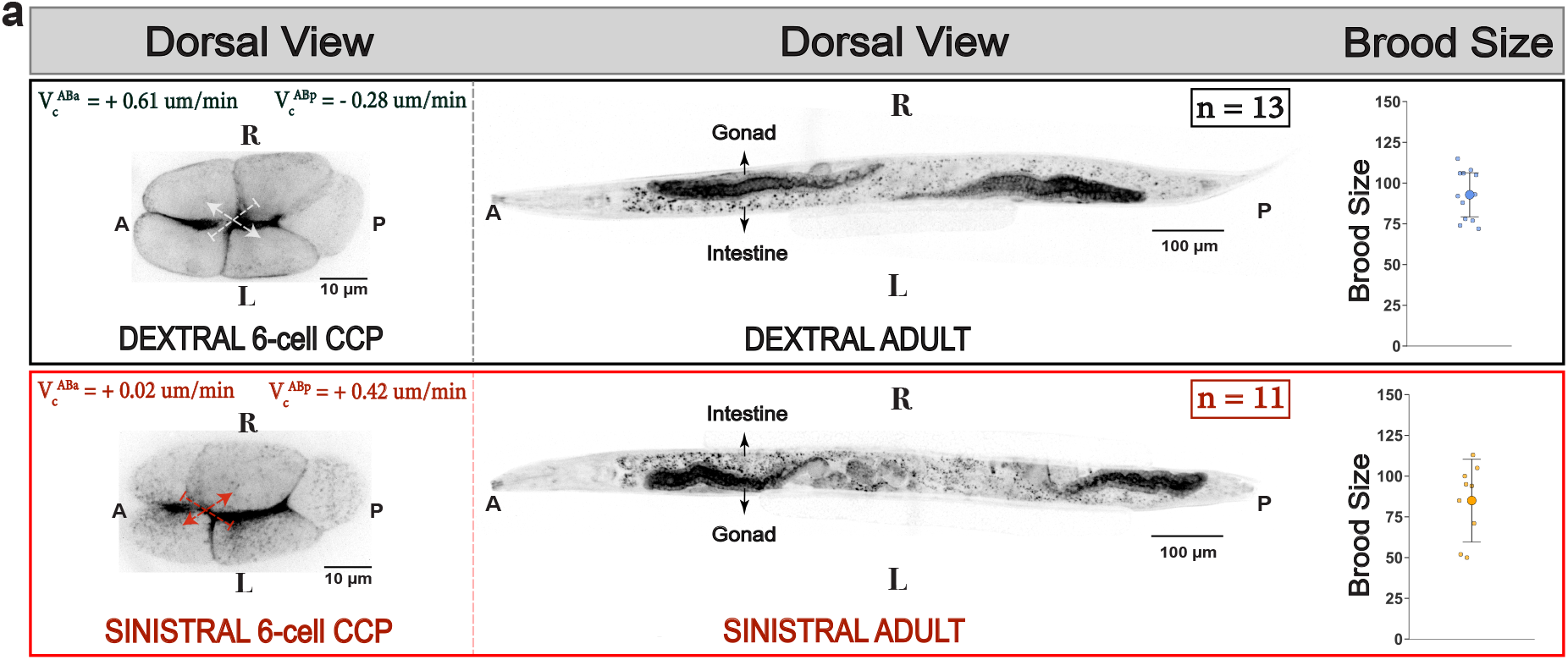
Effect of high amounts of Lifeact::mKate2 on LR handedness of C. elegans hermaphrodites. (a) 6-cell stage embryos and the corresponding adults they developed into, imaged from the dorsal side. Upper panel; a 6-cell embryo with dextral CCP, and the corresponding adult with dextral LR handedness. The respective 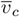 during the ABa and ABp divisions are reported for the representative embryo. The dorsal gonad should be on the right, and the intestine on the left of the adult hermaphrodite, to account for the stereotypic, or dextral LR handedness in C. elegans ([1][21][40] ; see Methods.) Upper right-most panel; brood size measured for such dextral adults (n=12) that developed from isolated 6-cell embryos with dextral CCP (for details on brood size measurement, see Methods). Lower panel; likewise for a 6-cell embryo with the sinistral CCP, and the adult that developed from it. The adult gonad-gut positioning is flipped when viewed from the dorsal side indicating sinistral LR handedness. Lower right-most panel; the brood size of sinistral adults (n=9) obtained from isolated 6-cell sinistral embryos. A two-sided Mann–Whiteney U Test was performed to verify that brood size is not significantly affected by the adult LR handedness (p = 0.48).

### Measurement of brood size

After imaging the adult for its gut-gonad chirality, the cover slip was gently removed and the immobilized adult was picked from the agar pad onto a new OP50 plate for further propagation. The immoblized worm recovered after ∼1 hr, and continued laying eggs. The adult, from this point, was moved to a new OP50 plate every 24 hrs until egg-laying stopped. All the OP50 plates, including the first one in which the worm had undergone its larval development were scored for hatched progeny, and summed to obtain the brood size. Fig.4a shows the brood size of the dextral adults compared to the sinistral adults.

### Particle Image Velocimetry (PIV) analysis

PIVlab(Matlab), was used to perform Particle Image Velocimetry (PIV) to compute 2D flow fields of the cortex [50]. A 3-pass filter with linear window deformation was used for PIV analysis. A final window size of 16x16 pixels with 8 pixel grid spacing was used for movies obtained with the 63x/1.2 NA(water) objective (pixel size = 0.1058 *µm*)[2][16]. For consistency, a 13 pixel grid spacing with a final window size of 26x26 pixels was used for movies obtained with the 100x/1.42 NA(oil) objective (pixel size = 0.0668 *µm*). PIV analysis was carried out on the maximum intensity projection of the z-stack.

Cortical flows in *C. elegans* zygotes were analysed from the onset of flow to pseudo-cleavage for ∼150 seconds. Note that the Lifeact++ zygotes under *rga-3(RNAi)* exhibited considerable ruffling of cell boundary during onset of polarity. To avoid the impact of ruffling on cortical flow fields, zygotes under *rga-3(RNAi)* were analysed for ∼ 100 seconds until ruffling was observed upon onset of cortical flow. The flow fields on *C. elegans* zygotes were rotated such that the A-P direction of the embryo aligned with the horizontal axis. In this orientation, the x-axis was along the long axis of the zygote, and the y-axis was orthogonal to it. The x-component *v*_*x*_ of the velocity vectors was parallel to the A-P direction, and positive when directed from anterior to posterior. The y-component *v*_*y*_ was perpendicular to the A-P direction, and positive when directed upwards from the long axis. Following alignment with the horizontal direction, the long axis of the embryo was spatially divided into 18 bins with a height of 120 pixels (12.7 *µm* for movies imaged with the 63x/1.2 NA objective), or 190 pixels (12.7 *µm* for movies imaged with the 100x/1.42 NA objective). *v*_*x*_ and *v*_*y*_ were spatially averaged in each bin, followed by a temporal average across the analysed frames to obtain the spatial profile along the A-P direction for a given embryo. The profiles shown in Fig.1b, f and Fig.1a,d are averages taken over embryos. The same procedure was used to calculate the mean NMY-2::GFP intensity profile along the A-P direction, shown in Fig.1a,d. Cortical flow fields during the AB division were analysed as in the time window stated in [16]. The AB cell was divided into 18 bins with a height of 94 pixels (9.94 *µm* for movies imaged with 63x/1.2 NA objective) or 154 pixels (9.88 *µm* for movies imaged with 100x/1.42 NA objective).

At the 4-6 cell stage, cortical flows were analysed for a period of 70-75 seconds from the onset of flows during cytokinesis until a 2.5 *µm* furrow depth during ingression was achieved by the ABa cell on its anterior tip, and by the ABp cell on its posterior tip. The ABa and the ABp cells were separately segmented for performing PIV analysis. The y-axis is parallel to the cytokinetic ring (along the A-P direction), and the x-axis is orthogonal to it (along the R-L direction). Therefore, x-component of a velocity vector *v*_*x*_ is positive when directed from right to left of the embryo, and the y-component *v*_*y*_ is positive when directed from anterior to posterior. The ABa cell was manually segmented and spatially divided into bins along the R-L direction with a height of 80 pixels (8.5 *µm*, 63x/1.2 NA objective). As previously stated, the flow field was spatially averaged in each bin, followed by a temporal average across the analysed frames to obtain the average profile for each embryo. The same was performed for the ABp cell separately. Figs.2c,g show average *v*_*y*_ profiles across the respective groups of embryos. Fig.2a shows the *v*_*y*_ profile from each individual embryo during the ABp division, along with average in each group. Fig. 3b shows the average *v*_*y*_ profile during ABa and ABp divisions separately for dextral and sinistral embryos.

### Measurement of 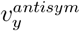 from average *v*_*y*_ profiles

For calculating the anti-symmetric *v*_*y*_ component 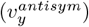 of cortical flows during the ABa division in a given 4-6 cell embryo, the average *v*_*y*_ profile obtained by PIV on the ABa cell(for more details, refer to the previous section) was reflected along the R-L direction and subtracted from the original unreflected profile. The average *v*_*y*_ profile for a given instance is an array of size (1,18) comprising of *v*_*y*_ values across 18 bins. These 18 bins are spatially positioned on the ABa cell along its R-L direction. Reflection of this array was obtained by a slicing operation with indexing [::-1]. The reflected array so obtained was subtracted from the original unreflected array (*v*_*y*_ - *v*_*y*_[::-1]) to obtain the antisymmetric component 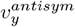 over the 18 bins. Likewise was done for the ABp cell, using average *v*_*y*_ profiles obtained from PIV performed during the ABp division in the ensemble of embryos. Fig. 3c shows the mean 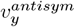 profile during the ABa division(left panel) and the ABp division (right panel). The profiles are averaged separately over ensembles of embryos that had dextral and sinistral CCPs at the 6-cell stage.

### Measurement of *v*_*c*_, 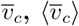

*v*_*c*_ for the *C. elegans* zygotes were calculated by averaging *v*_*y*_ on the posterior (across bins 13 to 16) and on the anterior (across bins 3 to 6) for each analysed frame, and subtracting the anterior average from the posterior average. For a given movie, we obtained a value of *v*_*c*_ corresponding to each analysed frame, and referred to it as the instantaneous chiral velocity [2, 12, 16, 51]. Fig.1c,g show distributions of *v*_*c*_ obtained from the observed zygotes in each respective group.

For the 4-6 cell stage, *v*_*c*_ was obtained by averaging *v*_*y*_ on the right (across bins 11 to 16) and on the left (across bins 2 to 7) of the ABa and ABp cells, and subtracting the right average from the left average for each cell. Please note that even though ABa and ABp divisions were always simultaneously imaged for a given embryo, data from the ABp cell was not analysed if the cortical plane drifted out of focus. Similarly, data from the ABa cell had to be discarded if the cell skewed out of focus during its division. Therefore, the ensemble of embryos used for calculation of *v*_*c*_ during ABa division is not identical to that used for the ABp division. Figs.2e,i and 2b,c show distributions and time evolution of *v*_*c*_ in ABa and ABp cells. Identical bins were used to define a posterior and an anterior region for calculation of *v*_*c*_ in the AB cell at the 2-3 cell stage (Fig. 1g).

Chiral velocity 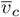 was obtained by temporally averaging *v*_*c*_ across all the analysed frames for each movie, both for the *C. elegans* zygotes and the 4-6 cell stage embryos [2]. Subsequently, the average chiral velocity 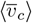 was obtained by averaging over 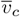 for a given ensemble of embryos.

### Intensity measurement Lifeact::mKate2 in *C. elegans* zygotes

Maximum intensity projections were used to segment the zygote. The area lying outside the zygote periphery was masked out, such that only the area lying inside had the original intensity values corresponding to Lifeact::mKate2 signal in the cortex. A single frame before onset of flow was selected such that the distribution of the Lifeact::mKate2 signal was uniform across the cortical plane. Intensity in the 594 nm channel used for imaging Lifeact::mKate2 in this frame was summed up in the segmented area to obtain the data for each embryo, shown in Figs.1h and 1d,f. Lifeact::mKate2 intensity values for zygotes of Lifeact-were assumed to be zero and included with its corresponding chiral velocities *v*_*c*_ to perform the linear fit with the data from the three Lifeact::mKate2 expressing transgenic strains. The intensites and chiral velocities were fit using the *’scipy*.*optimize*.*curve fit’* module of Python.

### Estimation of chirality index *c*

The active chiral fluid theory for the actomyosin cortex was used to estimate the chirality index *c*, as has been previously done [2][20][12][16]. Briefly, dynamics of the cortex are described by the following hydrodynamic equations:

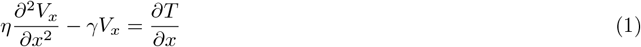

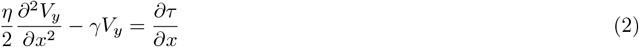

*η* denotes the viscous co-efficient of the cortex, and indirectly accounts for actin turnover at timescales longer than the typical turnover time of actin (∼ 30*s*, [51]). *γ* denotes the drag co-efficient experienced by the cortical flows from the surrounding medium (membrane/cytoplasm, [2]). *T* and *τ*, respectively denote the active tension and the active torque density in the cortex that have been shown to be driven by myosin activity [2][20][19]. Transforming to *x*^*′*^ = *x/L*, such that *L* is the length of the zygote along the A-P direction, and equating *T* and *τ* to linear functions of myosin intensity *I*, we obtain:

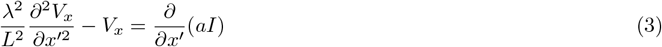

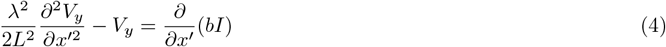

*λ* defined as 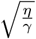 is the hydrodynamic length that denotes the length scale of cortical flow dissipation in response to a myosin gradient [52]. *a* and *b* are phenomenological constants that define *T* and *τ* as functions of myosin intensity *I*. The parameter chirality index *c* can be obtained from the phenomenological constants *a* and *b*, as follows [2][20]:

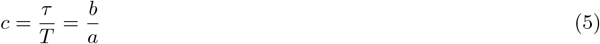

Using the myosin intensity profiles in Fig.1a,b, we can substitute *T* and *τ*, and numerically solve the constitutive hydrodynamic equations to obtain solutions for *v*_*x*_ and *v*_*y*_. The parameter set [*λ*, a, b] for the numerical solutions can be obtained by performing a least-square fit to the experimentally observed *v*_*x*_ and *v*_*y*_ profiles [2][51][16]. In our fittings, we kept the hydrodynamic length constant (*λ*=18 *µm*; 22 *µm* with *rga-3(RNAi)* [51]; with 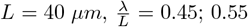 with *rga-3(RNAi)*), and used the least-squares fitting for estimating the parameters [a,b] to calculate *c*. L denotes the length of the long axis of the zygote, and 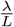 is referred to as the hydrodynamic ratio. 100 bootstrap iterations were used to obtain the average and 95% confidence to the mean. For each bootstrap iteration, a random collection of embryos, equal to the observed sample size, was drawn with repetition to obtain the average *v*_*x*_, *v*_*y*_ and myosin intensity *I* profiles, which were then used to calculate *c*. Specifically for the ABa and ABp cells, we made use of the average *v*_*x*_, 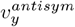 and myosin intensity profiles for calculating the chirality index, as the average 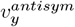 captured the rotational asymmetry in cortical flows better than the average *v*_*y*_ profiles. The hydrodynamic ratio was held constant at 0.45 for both the ABa and the ABp cells.

### Probability Measurements

For Fig.3, we classified the embryos with dextral and sinistral CCP at the 6-cell stage by the sign of their chiral velocity 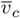 . We counted the number of embryos that had the dextral CCP and those that had the sinistral CCP at the 6-cell stage separately, and divided each by the total number of embryos in each group to calculate the respective probabilities. The error bars denote 95% confidence intervals (Clopper-Pearson exact CI), obtained via the *scipy*.*stats,beta*.*ppf* function in Python. For Fig.3b, the observed range of 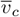 was divided into 0.5 *µm/min* bins, and probabilities for the dextral and the sinistral CCP at the 6-cell stage were measured in each bin.

### Cumulative Distribution Function (CDF) Calculation

The range of chiral velocities 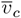 for 6-cell stage dextral and sinistral CCPs were divided into 0.5 *µm/min* bins. The probability distribution function (PDF) for a given sample of 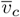 was analytically calculated using the following rule:

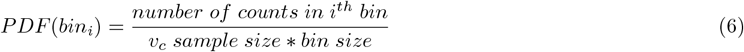

To smoothen the analytical PDF, the 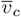 sample size was scaled in accordance with the PDF in each bin, and fit to a normal distribution using the python package ‘stats.norm.fit’. The mean and standard deviation obtained via the fitting were used to calculate the cumulative distribution function (CDF) using the ‘stats.norm.cdf’ package.

## Supplementary Information

### Movie legends

**S1**: *C. elegans* zygote exhibiting cortical flows with opposite handedness during onset of polarity. Upper panel; a maximum intensity projection of the cortex during polarity onset in a Lifeact-zygote expressing only NMY-II::GFP (left) and in a Lifeact++ zygote expressing both y NMY-II::GFP and Lifeact::mKate2 (right). The NMY-II::GFP signal is shown in gray; Lifeact::mKate2 in magenta. Anterior of the embryo is oriented to the left, posterior to the right. The maximum intensity projections were taken over 3 z-slices at the cortex with a separation distance of 0.5 *µm* between each slice. Bottom panel; the cortical flow profiles obtained from the NMY-II::GFP signal, overlaid with its maximum intensity projection for each frame. The last frame shows the temporally-averaged cortical flow profile over all the frames. Velocity vectors have the identical scale bar as shown in Fig.1a.

**S2**: Rescue of cortical flow handedness with suppression of Lifeact::mKate2 expression. Upper panel; a maximum intensity projection of the cortex during polarity onset in a Lifeact++ zygote (left) and in a Lifeact++ zygote under 24 hrs of *mKate2(RNAi)* (right). The NMY-II::GFP signal is shown in gray; Lifeact::mKate2 in magenta. Anterior of the embryo is oriented to the left, posterior to the right. The maximum intensity projections were taken over 3 z-slices at the cortex with a separation distance of 0.5 *µm* between each slice. Bottom panel; the cortical flow profiles obtained from the NMY-II::GFP signal, overlaid with its maximum intensity projection for each frame. The last frame shows the temporally-averaged cortical flow profile over all the frames. Velocity vectors have the identical scale bar as shown in Fig.1a.

**S3**: Reversal of cortical flow handedness amplifies upon hyper-activation of Rho-GTPase. Maximum intensity projections of the cortex in a Lifeact-zygote (left panel) and in a Lifeact++ zygote (right panel), respectively under 24 hrs and 12 hrs of *rga-3(RNAi)*. The NMY-II::GFP signal is shown in gray; Lifeact::mKate2 in magenta. Anterior of the embryo is oriented to the left, posterior to the right. The maximum intensity projections were taken over 3 z-slices at the cortex with a separation distance of 0.5 *µm* between each slice.

**S4**: Cortical flows during onset of polarity in zygotes of other transgenic worms strains expressing Lifeact::mKate2. Maximum intensity projections of the cortex in a SWG050 zygote expressing only Lifeact::mkate2 (left) and in a SWG218 zygote expressing both NMY-II::GFP and Lifeact::mKate2 (right). The NMY-II::GFP signal is shown in gray; Lifeact::mKate2 in magenta. Anterior of the embryo is oriented to the left, posterior to the right. The maximum intensity projections were taken over 3 z-slices at the cortex with a separation distance of 0.5 *µm* between each slice. For more details about the transgenic constructs, see Methods.

**S5**: Cortical flows during the 4-6 cell stage in a Lifeact-embryo, imaged from the dorsal side (upper panel). Anterior of the embryo is oriented to the left, posterior to the right. Movie shows maximum intensity projection of the NMY-II::GFP signal over 23 z-slices with a separation distance of 0.5 *µm* between each slice; starting at the cortex on the dorsal side and progressing inwards into the embryo. Bottom panel; the cortical flow profiles obtained from the NMY-II::GFP signal on the ABa cell, overlaid with its maximum intensity projection for each frame. The last frame shows the temporally-averaged cortical flow profile over all the frames. Velocity vectors have the identical scale bar as shown in Fig.2b.

**S6**: Cortical flows during the 4-6 cell stage in a Lifeact++ embryo, imaged from the dorsal side (upper panel). Anterior of the embryo is oriented to the left, posterior to the right. Movie shows maximum intensity projection of the NMY-II::GFP signal over 15 z-slices with a separation distance of 0.75 *µm* between each slice; starting at the cortex on the dorsal side and progressing inwards into the embryo. Bottom panel; the cortical flow profiles obtained from the NMY-II::GFP signal on the ABa cell, overlaid with its maximum intensity projection for each frame. The last frame shows the temporally-averaged cortical flow profile over all the frames. Velocity vectors have the identical scale bar as shown in Fig.2b.

**S7**: Cortical flows during the 4-6 cell stage in a Lifeact-embryo, imaged from the dorsal side (upper panel). Anterior of the embryo is oriented to the left, posterior to the right. Movie shows maximum intensity projection of the NMY-II::GFP signal over 23 z-slices with a separation distance of 0.5 *µm* between each slice; starting at the cortex on the dorsal side and progressing inwards into the embryo. Bottom panel; the cortical flow profiles obtained from the NMY-II::GFP signal on the ABp cell, overlaid with its maximum intensity projection for each frame. The last frame shows the temporally-averaged cortical flow profile over all the frames. Velocity vectors have the identical scale bar as shown in Fig.2f.

**S8**: Cortical flows during the 4-6 cell stage in Lifeact++ embryos imaged from the dorsal side, classified by direction of cortical counter-rotation during the ABp division. Right panel; a Lifeact++ embryo showing stereotypic directionality of cortical counter-rotational flows during the ABp division, identical to the Lifeact-zygotes; observed in 21/38 embryos of Lifeact++. Center; a Lifeact++ embryo showing reversed directionality of cortical counter-rotational flows during the ABp division, observed in 10/38 embryos of Lifeact++. Left panel; a Lifeact++ embryo showing cortical flows directed towards the anterior of the embryo during the ABp division, observed in 7/38 embryos of Lifeact++. Anterior of each embryo is oriented to the left, posterior to the right. Movie shows maximum intensity projections of the NMY-II::GFP signal over 15-20 z-slices with a separation distance of 0.75 *µm* between each slice; starting at the cortex on the dorsal side and progressing inwards into the embryo. Bottom panel; the cortical flow profiles obtained from the NMY-II::GFP signal on the ABp cell, overlaid with its maximum intensity projection for each frame. The last frame shows the temporally-averaged cortical flow profile over all the frames. Velocity vectors have the identical scale bar as shown in Fig.2f.

**S9**: Representative Lifeact++ embryos undergoing ABa and ABp cell divisions to give rise to the 6-cell stage cell-cell contact pattern (CCP). From right to left; the dextral CCP, the indeterminate CCP, the sinistral CCP. The arrows overlaid on the last frame indicate the respective 6-cell stage CCPs. Pointed arrow heads denote the cells in contact; flat arrow heads denote the cells not in contact. Anterior of each embryo is oriented to the left, posterior to the right. Movie shows maximum intensity projections of the NMY-II::GFP signal over 15-20 z-slices with a separation distance of 0.75 *µm* between each slice; starting at the cortex on the dorsal side and progressing inwards into the embryo.

**S10**: Z-stacks obtained from Lifeact++ hermaphrodites oriented ventral side-up, with the dextral body plan (upper panel) and the sinistral body plan (bottom panel). Movies show 52 z-slices 1 *µm* apart, starting on the ventral surface of the worm and progressing inwards into the body. NMY-II::GFP in gray; Lifeact::mKate2 in magenta.

## Acknowledgments

We thank Teije Middelkoop, Jonas Neipel, Archit Bhatnagar, Alexandra Schauer and Amin Tajik for fruitful discussions, and Ella Linxia Müller for inputs on mounting and imaging of the 4-6 cell embryo.

